# Genome-wide association mapping identifies Yellow Rust resistance locus in Ethiopian Durum Wheat germplasm

**DOI:** 10.1101/2020.11.27.400895

**Authors:** Sisay Kidane Alemu, Ayele Badebo Huluka, Kassahun Tesfaye Geletu, Cristobal Uauy

**Affiliations:** National Agricultural Biotechnology Research Center, Holeta, Ethiopian Institute of Agricultural Research, Addis Ababa, Ethiopia; International Maize and Wheat Improvement Center (CIMMYT), Addis Ababa, Ethiopia; Institute of Biotechnology and DMCMB, Addis Ababa University; Addis Ababa, Ethiopia; John Innes Centre, Norwich Research Park, NR4 7UH, Norwich, UK

**Keywords:** GWAS, Association, SNP, QTLs, Stripe rust, Coefficient of Infection, Field Resistance

## Abstract

Durum wheat is an important cereal grown in Ethiopia, a country which is also its center for genetic diversity. Yellow (stripe) rust caused by *Puccinia striiformis* fsp *tritici* is one of the most devastating diseases threatening Ethiopian wheat production. To identify sources of genetic resistance to combat this pathogen, we conducted a genome wide association study of yellow rust resistance on 300 durum wheat accessions comprising 261 landraces and 39 cultivars. The accessions were evaluated for their field resistance in an alpha lattice design (10 × 30) in two replications at Meraro, Kulumsa and Chefe-Donsa in the 2015 and 2016 main growing seasons. Disease Scoring was carried out using a modified Cobb scale and then converted to Coefficient of Infection (CI). Analysis of the 35K Axiom Array genotyping data resulted in a total of 8,797 polymorphic SNPs of which 7,093 were used in subsequent analyses. Population structure analysis suggested two groups in which the cultivars clearly stood out separately from the landraces. We identified twelve SNPs significantly associated with yellow rust resistance across four chromosomes (1A, 1B, 2B, and 7B). Six of the SNPs (AX-95171339, AX-94436448, AX-95238778, AX-95096041, AX-94730403 & AX-94427201), were consistently identified on chromosome 1B at the three field locations and combined across the six environments. The phenotypic variation (R^2^) explained by all six SNPs on chromosome 1B ranged from 63.7 – 65.4%. Locus-based analysis of phenotypic values between resistant and susceptible allele resulted in a significant difference at (p < 0.001). Further investigation across the genomic interval encompassing the identified loci indicated the presence of disease resistance protein (NBS-LRR class) family and RPM1 in the vicinity of the loci. This study provides SNPs for tracking the QTL associated with yellow rust resistance in durum wheat improvement programs.

## Introduction

Durum wheat (*Triticum turgudum* subsp. *durum*) is a tetraploid wheat (2n=4x=28) with AABB genome designation and basic chromosome number x=7. Tetraploid wheat is thought to be the result of a genome hybridization between the diploid A genome wheat species *Triticum urartu* (2n=2x=14) and an S genome species related to *Aegilops speltoides* [1].

Durum wheat is an important food crop with global production estimated to be 36 million t per year [2].Ethiopia is considered as a center of secondary diversity for durum wheat and as such has a wide array of untapped tetraploid germplasm [3–9]. Although statistics on current production are combined (in most cases) into a single “wheat” category, occasional studies report that Ethiopia is the largest producer of durum wheat in sub-Saharan Africa with approximately 0.6 million hectares (Evan School Policy Analysis and Research [10]. Particularly, landraces (i.e. a locally adapted line that has not been modified through a breeding programme) are known to cover about 70% of the total durum wheat area [11]. The crop is grown widely on heavy black soils of the highlands with an altitude range of 1800 to 2800 masl [12, 13]. Due to this continuous production by farmers in the highlands, Ethiopian durum wheat has evolved in its phenotypic [14] and molecular diversity [15]. Selection pressure from the natural and artificial sources, inevitably has contributed to the development of many adaptive traits including disease resistance [16–18] which provide an opportunity for genetic improvement programs.

Durum wheat is mostly grown as a food crop by small holder farmers, although it has potential as an industrial crop in the food industries. Pasta, spaghetti, biscuits, pancakes, macaroni, pastries, and unleavened breads are among the known forms of uses of durum wheat [19]; Global durum wheat use trending upward [20]. Other traditionally used Ethiopian recipe includes dabo (home-made bread), ambasha (bread from northern Ethiopia), kitta (unleavened bread), nifro (boiled whole grains), kolo (roasted whole grains), dabo-kolo (round and seasoned dough) and kinche (crushed kernels, cooked with milk or water and mixed with spiced butter) [21]. In the mixed farming systems of the highlands, the straw of durum wheat is also of high relevance for animal feed due to its digestibility and palatability characters [22].

Stripe/yellow rust of wheat is a fungal disease caused b *y Puccinia striiformis* f. sp *tritici* (Pst), an obligate biotrophic fungal pathogen which depends on living cells of the host for its survival and reproduction [23]. It has both sexual and asexual stages and it is the asexual stage which is pathogenic and damages wheat through the infectious units called urediniospores [23]. Stripe rust is one of the most devastating diseases in wheat growing regions of Ethiopia and it has been considered as a regular disease, occurring in all cropping seasons [24]. In 2010, an epidemic of the disease occurred in the country and reached all the wheat growing regions at unprecedented rates. It infected large production areas including 29 zones in the Amhara, Oromia, and Southern Nations, Nationalities, and Peoples (SNNP) regional states [25–26]; http://www.addisfortune.com.

Stripe rust can be managed using various methods such as cultural practices, fungicide application, and resistance cultivar development and deployment [27–30], with resistant variety deployment being an environmentally friendly strategy. Due to the dynamic nature of the pathogen, new virulent Pst races often emerge and break varietal resistance leading to disqualification from production of cultivars. This situation prompts the necessity of continuous search and identification of new sources of resistance to sustain wheat production.

In Ethiopia, research effort made by the wheat improvement programs has resulted in the development of cultivars with relatively good levels of wheat rust resistance, good yield, and agronomic characters [31]. Germplasm screening to identify resistant breeding lines and development of elite durum wheat cultivars [32] has been part of the strategies to exploit the host’s natural defence in breeding programs. Research reports on wheat rusts so far indicate that about 82 *Yr* genes have been identified which are known to confer both adult plant resistance (APR) and all stage (seedling resistance) [33–35]. Screening available germplasms for the presence of these known genes is a strategy to maximize the resistance potential of that germplasm. Specifically, when the genes are mapped to a known genomic locus and diagnostic molecular markers are available, the detection of the known resistant genes is facilitated. However, several of the identified genes have limited spectra of effectiveness as they have been previously overcome by specific Pst races. Besides, germplasm pools vary in their genetic background and local adaptation making it difficult in the absence of perfect markers to decipher which known *Yr* resistance genes they carry. A strategy to address this is searching for resistance genes in locally adapted germplasm using techniques such as Genome Wide Association Studies (GWAS).

GWAS is is a powerful study design to identify genetic variants of a trait through detection of the association between a single-nucleotide polymorphisms (SNPs) and the trait of interest [36–37]. GWAS has been used extensively in plant studies since the beginning of this century [38]. Following earlier reports of GWAS related work on maize [39] and *Arabidopsis* [40], several reports began to be published in other crops as reviewed in Rafalski [41]. There are also reports from GWAS studies on identification of the genetic basis of farmers preferences on durum wheat traits (Kidane et al., 2017b [42]). The availability of a well characterized population, usually called an association or diversity panel, is a pivotal requirement to conduct such studies. The panel is genotyped with an appropriate platform (often SNP genotyping) and phenotyped for the trait of interest in a designed trial to generate the molecular and trait data. Once phenotype and genotype data are available, the remaining task is to run the association analysis using dedicated computer programs and appropriate statistical models [37, 43]. GWAS studies ultimately result in the identification of genomic loci/SNPs which are significantly associated with the target traits [37] and can be used in marker assisted introgression programs to improve the target crop. As a follow-up investigation, the associated SNPs can be validated for their diagnostic values in an independent germplasm.

Ethiopian durum landraces have been suggested as a good source of resistance to wheat rusts [44–45]. This has been supported by association studies on relatively small sized Ethiopian durum panels in limited study environments [17]. More extensive research on potential new and effective Pst resistance associated loci or genes is necessary to cope with the occurrence of new virulent *Pst* races. The availability of high-density SNP wheat chips e.g. 35k Breeders chips, iSelect 9 and 90 K SNP assays [46–48] provides an opportunity to facilitate such genome level studies. The objectives of this study were: (1) to access the extent of field resistance to stripe rust among an Ethiopian durum wheat panel and (2) to identify significantly associated SNP markers with the resistance genomic loci.

## Materials and Methods

### Phenotyping

#### Plant Materials

Durum wheat accessions (n=513) were obtained from various sources in Ethiopia. The accessions comprise landraces from gene bank collections (Ethiopian Biodiversity Institute (EBI) and Ethio-Organic Seed Action (EOSA)) and landraces and released varieties from Debre-Zeit Agricultural Research Center (DZARC). To start with a relatively pure seed stock, all the accessions were grown and those having a mixed seed stock were identified by physical observation on morphological features such as spike architecture (e.g. density, color, awns) and seed color. In few instances, accessions of mixed genotypes were split and considered as a sub accession making the final number higher. Each accession was then subjected to a single spike row planting followed by two generations of self-pollination. A final single spike to row planting was carried out at Holeta Agricultural Research Center for seed multiplication. The final working population was then constituted and cut down to 300 based on morphological similarity among the accessions and used for phenotypic evaluations and genotyping. This final panel included 261 landraces and 39 cultivars. The source and related description of the accessions is provided in S1 Table.

#### Field Resistance Evaluation

The accessions were grown in an alpha lattice design with two replications at Chefe-Donsa (CHD), Kulumsa (KUL) and Meraro (MER) sites in the main growing season (June – November) of 2015 and 2016. Chefe-Donsa is located 35 kms east of Debre Zeit at 08°57’15” N and 39°06’04” E and has an altitude of 2450 m. Kulumsa is located 167 kms from Addis Ababa at 8°01’11.7"N 39°09’38.2″E and has an attitude of 2200 m; whereas Meraro is located about 236 kms from Addis Ababa at 7°24’25.8″N 39°14’56.3″E and has an elevation about 3,030 m. Each accession in each replication was sown in two rows of 0.5 m length, with 0.2 m spacing between rows. Each block was enclosed between spreader rows of known susceptible durum wheat (LD-357, Arendato and Local Red) and bread wheat (Morocco known to have Sr25 and Lr19) varieties mixed in the ratio of 1:1:1:1 and sown 20 cm from the experimental plots. The spreader rows act as an inoculum source and help in achieving uniform disease establishment throughout the experimental field. The disease severity (percentage of leaf tissue infected with the rust) was evaluated using a modified Cobb’s scale [49] with values ranging from 0 to 100%. The field response of the genotypes to the rust infection was scored according to Stubbs et al.

[50] as R (resistant), MR (Moderately Resistant), Moderate (M), MS (Moderately, Susceptible) and S (Susceptible) each having a numerical constant value of 0.2, 0.4, 0.6, 0.8 and 1.0, respectively. All agronomic practices were applied following the recommended practice for wheat at each location. Two data sets (one for each year) per each of the three locations was generated giving a total of six environments.

#### Phenotype Data Analysis

Severity (SEV) score was multiplied with the field Response (RES) values to produce the Coefficient of Infection (CI) which represents the overall reaction of the genotypes for the pathogen. The panel was classified in to Resistant, Intermediate and Susceptible groups based on the average values of SEV and RES [17, 51] for each environment. As CI is the product of SEV and RES, the corresponding values were used to do the same reaction classification in terms of CI as well. To comply with the normality assumption, SEV and CI data were transformed with common logarithmic function (SEV_tr_ = LOG_10_(SEV+1) and CI_tr_ = LOG_10_(CI+1)) while Response was subjected to arcsine transformation (RES_tr_ = arcsine(√RES)). Shapiro Wilk test (Shapiro and Wilk 1965 [52]) was applied on the original and the transformed data to assess the normality of the data. Analysis of Variance (ANOVA) was done with the transformed data for each and combined over environment using a linear mixed model. In the model, genotype, Replication, Incomplete block within Replication, Location, year, and genotype by other variance source interactions were considered as random effects. Multiple random effect test (Variance component test) was carried out using Lym4 and LimerTest packages in R. The Best Linear Unbiased Estimates (BLUEs) of each environment and combined data (BLUE-all) was generated using Restricted Maximum Likelihood (REML) method fitting the genotype as fixed effect while the other variance sources and interactions as random effect. These BLUE values were used to perform the association analysis. Correlation analysis was performed for SEV, RES and CI to assess the extent of covariation of the resistance reaction within and across environments.

### Genotyping

Seeds were sown on a 50 mm diameter sterile Petri dish equipped with 42 mm filter discs to maintain a moist condition for germination. After watering them with distilled water, Petri dishes were incubated at 4°C under dark conditions for 24 hours to break dormancy. The dishes were then kept at room temperature for 3-4 days until fully germinated. Germinated seeds were planted in a 96 well tray filled with peat and sand soil mix optimized for raising cereal seedlings. The trays were placed in cereal growth chamber set at 19°C day and 16°C night temperature with a relative humidity of 70% and a photoperiod of 16:8 hours light/dark cycle.

Fully opened leaves were harvested from 10 to 14 days old seedlings in a 1.2 mL deep-well plate on a dry ice and freeze-dried for 24-30 hours at −40°C under a pressure of 20 atm. The tissue was then ground into a fine powder followed by a wet grinding with Geno/Grinder 2010 at 1750 rpm for 2 minutes. Genomic DNA was extracted with SDS buffer following the wheat and barley DNA extraction protocol in 96-well Plates [53] with some modification. DNA was cleaned according to Affymetrix User Guide, Axiom^®^ 2.0 Assay for 384 Samples (Genomic DNA Preparation and Requirements) and quantified with Nanodrop (8-sample spectrophotometer ND-8000). A total of 100 μL of DNA sample normalized to 75-100 ng/μL was submitted to University of Bristol Genomic Facility for genotyping using Breeders’ 35K Axiom Arra. At the service center, sample array processing and genotyping was carried out following the Axiom^®^ 2.0 Assay for 384 Samples user and Workflow guide (http://media.affymetrix.com/support/downloads/manuals/axiom_2_assay_auto_workflow_user_guide.pdf). We received the genotype data as ARR, JPEG and CEL files, where the latter was used in downstream SNP and association analyses.

### Analysis

#### SNP/Genotype Data Analysis

Genotype/SNP analysis was performed with Axiom Analysis Suit (AxAS) v2.0 using the CEL intensity files following sample QC and the Best Practice Workflow. Poor quality samples were identified with Dish Quality Control (DQC) values where samples having a value of ≤ 0.80 were excluded from the next step of the analysis. Using a subset of probe sets, samples which passed the DQC value step were subjected to genotype calling to generate the QC call rate and those samples which had a value ≤ 0.91% were excluded from all subsequent genotyping and SNP data analysis. Once the good quality (QC passed) samples were identified, the downstream genotype data analysis was performed following the Best Practice Workflow with SNP QC default settings for SNP genotype calling. The resulting SNP classes were assessed for complying with expected thresholds mainly with the number of minor alleles ≥ 2. Accordingly, the ‘polyhighresolution’ and ‘NoMinorHomos’ classes, as they fulfill the criteria, were closely examined, and considered for the next step analysis. The other SNP classes (MonoHighResolution, OTV, Other and CallRateBelowThreshold) were not considered because they are noisy in many aspects and did not comply with the minor allele number threshold mentioned above.

The SNP summary table and the genotype call data were extracted using the export tab of the Axiom analysis window. The accessions’ genotype data was further subjected to individual heterozygosity analysis and accessions with a value of ≥ 3% heterozygosity were excluded from the panel. The physical positions of the SNPs were extracted from the position file of Breeders’ 35K Axiom Array anchored on the wheat reference genome sequence RefSeq v1.0 [54]. Minor Allele Frequency (MAF) was calculated as a percentage of each SNP allele relative to the total in the association panel and individuals having a MAF value > 5% (cut-off) were considered for the downstream analyses. Furthermore, Locus Heterozygosity (the probability that an individual is heterozygous for the locus in the population and polymorphism Information content (PIC: the discriminating power of the marker in a population) were calculated as describe in Liu [55–56]

#### Population Structure & Kinship

Population structure among the association panel was analyzed using STRUCTURE v 2.3.4. [57]. Allele frequency model was employed at Length of burnin period: 10000, number of MCMC Reps after burnin: 100000, K runs from 1 - 10 with 5 times replication for each K. The result of the analysis was zipped in to a folder and uploaded on to STRUCTURE HARVESTER [58], an online analysis tool used to generate an estimate of the optimum number of subpopulations (i.e. the value of K) through Evanno method [59]. Kinship matrix was generated through genetic similarity matching using all possible pairwise combination among the panel using the R package Genomic Association and Prediction Integrated Tool (GAPIT) besides performing the PCA [60].

#### Linkage Disequilibrium

Genome-wide Linkage Disequilibrium was assessed using TASSEL v.5 to estimate the squared allele frequency correlation (r^2^) for all pairwise comparison of distances between SNPs. The genotype data was imported into TASSEL in HAPMAP format and full matrix LD analysis was performed with the default settings. The resulting LD output was subjected to binning using customized R script with bin_size of 1000 bp and maximum_bin of 829100000 bp to generate the average r^2^ and the inter-SNP distance values. The LD values were plotted against the corresponding inter-SNP physical distance using a customized R script to estimate the LD decay rate as described in Hill and Weir [61]. The threshold value of the LD (r^2^) was set at 0.2 a commonly applied value in many related studies [62–65. To visualize the LD decay pattern, nonlinear model was fitted to the LD plot relating the squared allele frequency with the physical distance [66].

#### Association Analysis and Locus Exploration

The working set 7093 SNP genotype data across the 293 genotypes was organized in the HapMap format. The phenotype data in terms of the Best Linier Estimate (BLUE) for SEV, RES, and CI was arranged across the 293 genotypes for all the single environment data sets. GAPIT [60] was used for the association analysis in R v 3.6.2 and RStudio v. 1.2.5033 to identify the genomic loci underlying the *Pst* resistance. The markers and the phenotype data corresponding to each genotype was input and fitted in the compressed mixed linear model (MLM) which accounts for uneven relatedness and controls effectively through lowering the type I errors [67]. VanRaden’s method [68] was applied to calculate Kinship. To account for the genetic structure, Principal component Analysis (PCA) was calculated and iteratively added to the model [69]. The best fit of the model was visually assessed by observing the QQ plots. Correction for false positive association was then performed by adding the default K + PCA covariates to the fixed effect part of the model in the GAPIT code. The analysis was carried out first at each environment using a single dataset, then combined across locations, years, and all environments. The probability of adjusted false discovery rate (p < 0.05) was used as a critical value to declare the significant marker trait association [70]. The “-log10(p)” values were plotted against each chromosome and the “expected −log10(p)” to generate Manhattan and QQ-plots respectively with codes imbedded in the GAPIT script. The SNP allele in the most and consistently susceptible line EDW-262 was used as the susceptible allele and the alternative was considered as the resistant one. The resistant allele frequency among the panel was determined based on the number of resistant alleles. Single locus-based variation of phenotypic values of resistant and susceptible alleles was also tested using Two-Sample T-test assuming equal variances. Further exploration was done on approximately 4.15 million (bp) genomic region encompassing all six identified loci on chromosome 1B. Genes and gene models reported with in the identified resistance associated region and close to the nearest significant SNP were examined using list of Genes/gene model extracted from the wheat genome annotation file [54].

## Results

### Phenotypic Reaction to Pst Infection

Variably distributed reaction groups were demonstrated among the panel for SEV, RES and CI across the six environments (Fig 1). For SEV, 100% of the accessions were classified as resistant (0≤SEV≤10) at CHD_15 and 99.7% at KUL_15 while 49, 95, 84, 58% were classified at MER_15, CHD_16, KUL_16 and MER_16 respectively (Fig 1A). For RES, 96.6% of the accessions fell under resistant class at CHD_15 and 97.63% at KUL_15, while 33, 91, 79, 41% were observed under the same resistant class at MER_15, CHD_16, KUL_16 and MER_16 in that order(Fig 1B). Almost similar frequency of reaction groups were observed for CI where 99% of them were under resistant class both at CHD_15 and KUL_15 while 38, 89, 78, 49% appeared in the same reaction group respectively at MER_15, CHD_16, KUL_16 and MER_16 (Fig 1C). The mean SEV, RES and CI values across the environments ranged from 0-63.8%, 0-0.83 and 0-63.8 in the same order. Considering stability of reactions across all the six environments, 37, 26, and 31% of the accessions gave consistently resistant reaction for SEV, RES and CI across the testing location in that order.

**Fig 1.**
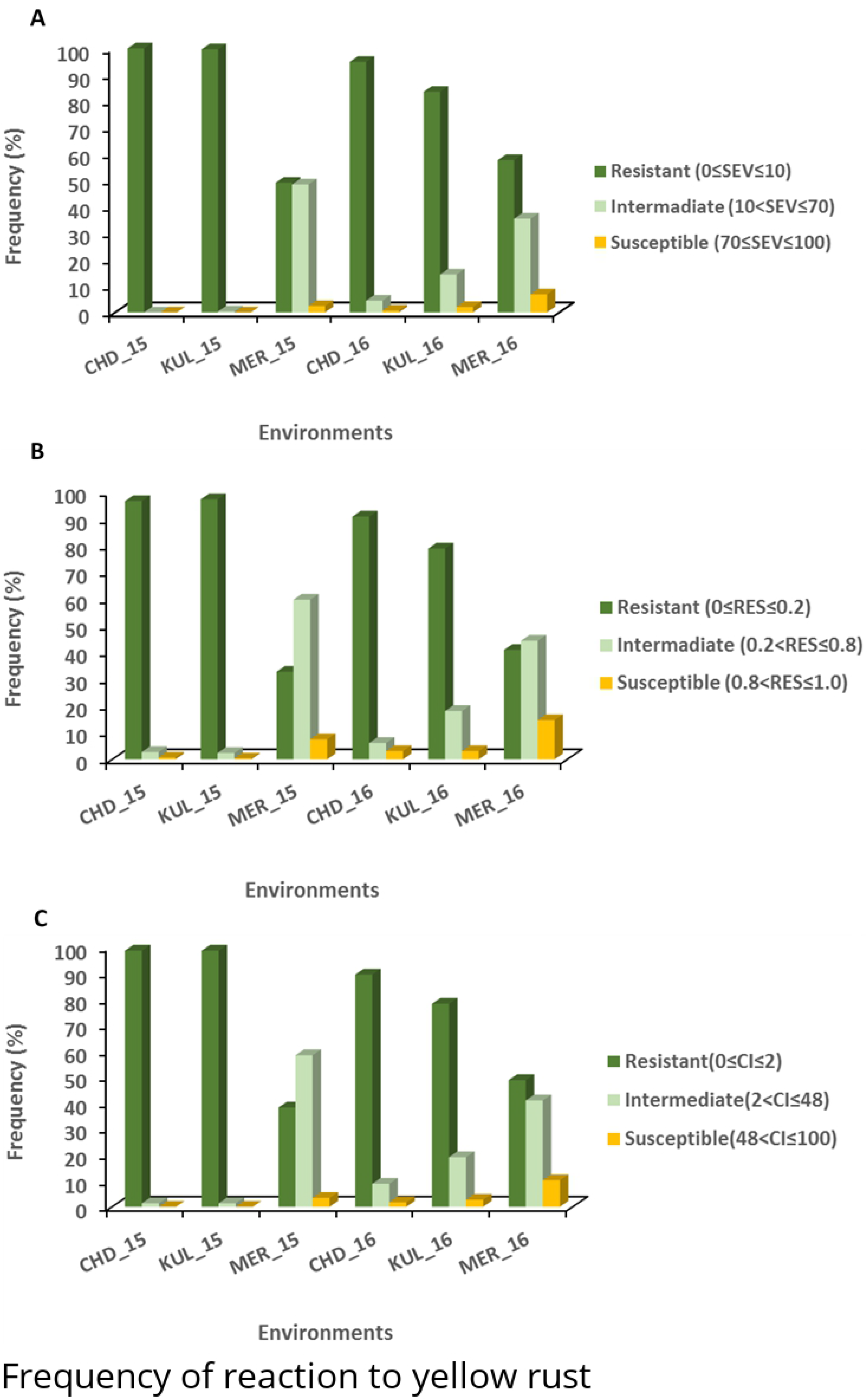
Reaction to yellow rust of 293 Ethiopian durum wheat accessions obtained from field experiments in six environments. A) severity, B) Field Response and C) Coefficient of Infection. The reaction data of severity and field response was used to classify the panel in to the three response groups as described in Liu et al., 2017 [17] for severity and Field Response and taking the corresponding values for Coefficient of Infection.

Regardless of the various reaction groups, the frequency distribution of the original untransformed data (BLUE-all-Original) values was heavily skewed for SEV (W=0.67) and CI (W=0.57), while Response appeared to be less skewed (W=0.89) (Fig 2A, B, C). Application of logarithmic transformation to SEV and CI and the arcsine transformation to RES changed the distribution to better adjust to normality with W values of 0.92 for SEV, 0.93 for RES and 0.87 for CI (Fig 2D, E, F).

**Fig 2.**
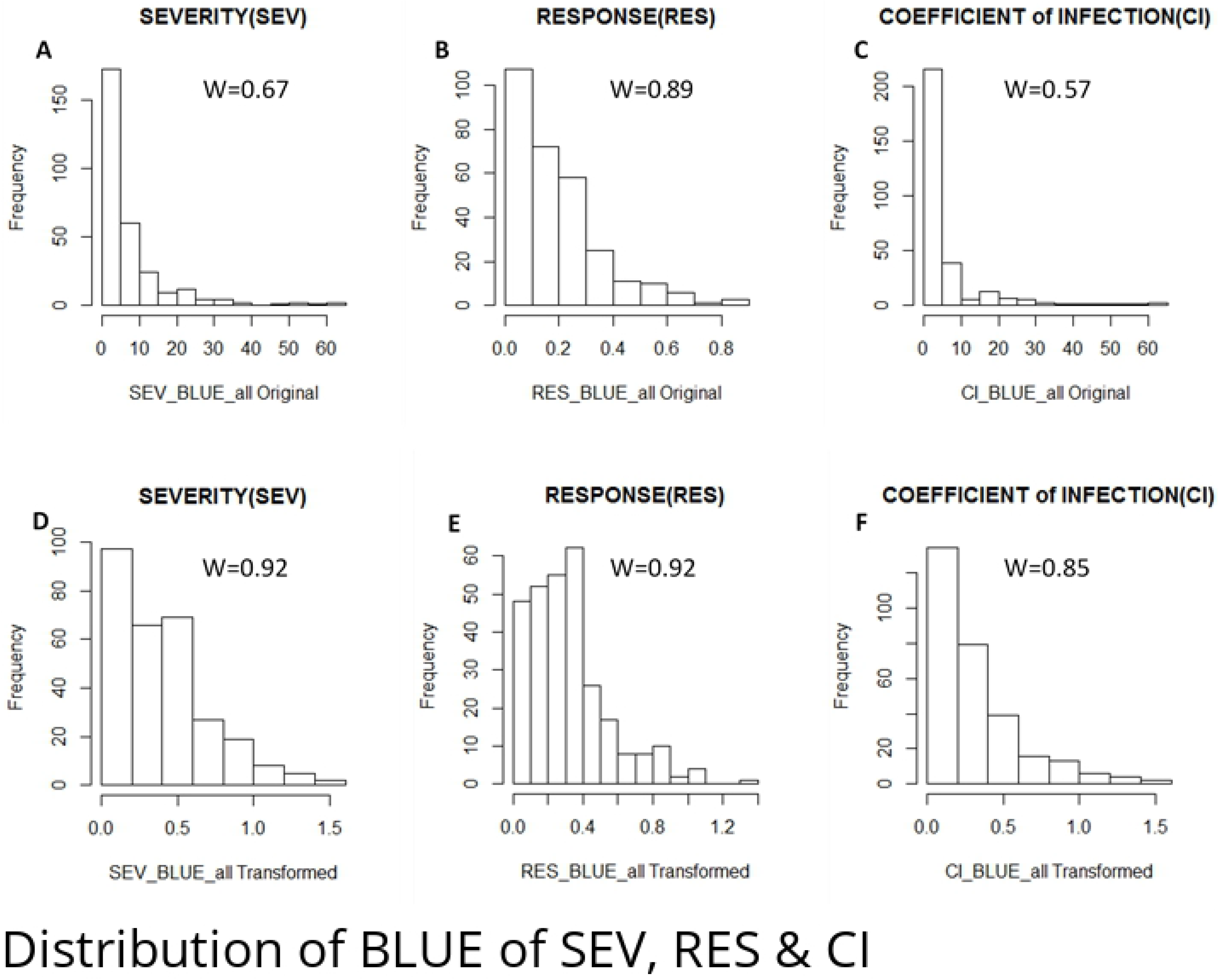
Distributions of disease severity (SEV), Response (RES) and Coefficient of Infection (CI). Distributions of best linear unbiased estimates (BLUEs) across six environments for SEV, RES and CI are represented by A, B and C for Original data while by D, E and F for Transformed data. “W” is Shapiro - Wilks Statistic indicating the correlation between observed values and normal scores both for the original and transformed values.

### Variances and Correlations of Reactions

We first examined the variance components for the multiple factors included in the statistical model. ANOVA of location and combined data across environments is summarized in Table 1. Genotypic variance components were significant for most of the test locations (*P < 0.001*) and combined across environments except for SEV of KUL (Table 1). Replication was significant (*P < 0.001)* at Kulumsa, Meraro and Chefe-Donsa (*P < 0.01*) while it was nonsignificant for the combined analysis. Blocks nested within replications were non-significant both for the individual locations and combined across locations. Variance components of locations and years were significant (*P < 0.001*) for the combined analysis and the individual locations except for MER. Genotype by environment interactions (two-way) were significant (*P < 0.001)* at the individual locations (GxY) and across the combined data (GXL & GxY) for all SEV,RES & CI. Likewise, the three-way interaction (GxLxY) resulted in a highly significant variation at *P < 0.001* for SEV, RES and CI. Apparently, this shows that there was strong interaction between the response of each genotype based on the location and year, consistent with the data presented in Fig 1.

**Table 1.**
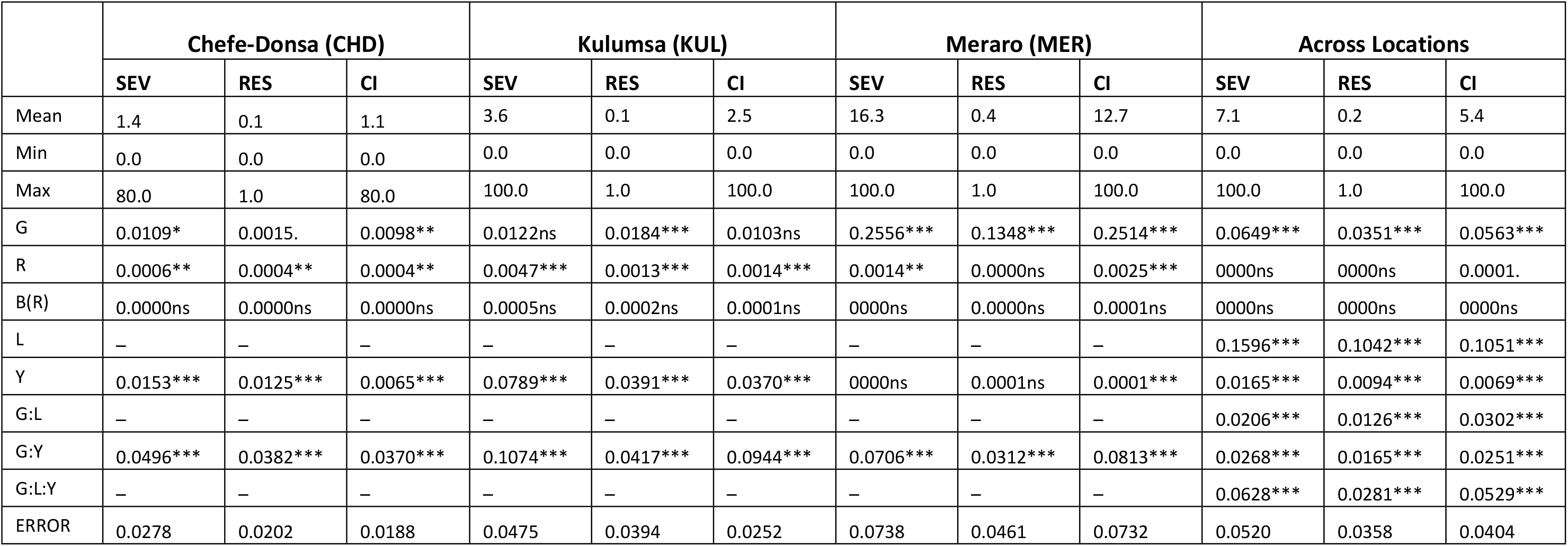
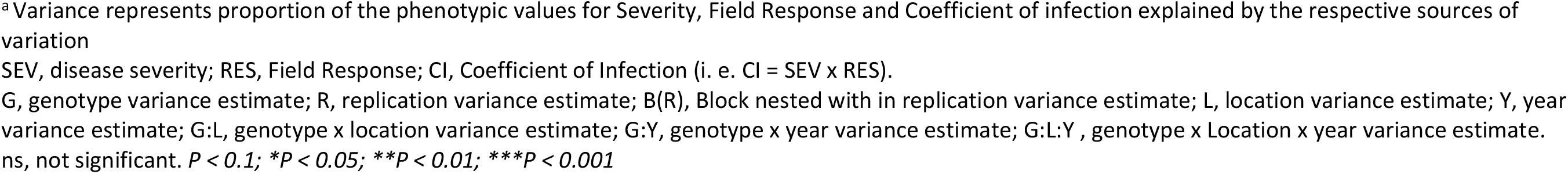
Variance^a^ in reaction to yellow rust of 293 Ethiopian durum wheat accessions per test location over two years and combined across environments.

|  | Chefe-Donsa (CHD) |  |  | Kulumsa (KUL) |  |  | Meraro (MER) |  |  | Across Locations |  |  |
| --- | --- | --- | --- | --- | --- | --- | --- | --- | --- | --- | --- | --- |
|  | SEV | RES | CI | SEV | RES | CI | SEV | RES | CI | SEV | RES | CI |
| Mean | 1.4 | 0.1 | 1.1 | 3.6 | 0.1 | 2.5 | 16.3 | 0.4 | 12.7 | 7.1 | 0.2 | 5.4 |
| Min | 0.0 | 0.0 | 0.0 | 0.0 | 0.0 | 0.0 | 0.0 | 0.0 | 0.0 | 0.0 | 0.0 | 0.0 |
| Max | 80.0 | 1.0 | 80.0 | 100.0 | 1.0 | 100.0 | 100.0 | 1.0 | 100.0 | 100.0 | 1.0 | 100.0 |
| G | 0.0109* | 0.0015. | 0.0098** | 0.0122ns | 0.0184*** | 0.0103ns | 0.2556*** | 0.1348*** | 0.2514*** | 0.0649*** | 0.0351*** | 0.0563*** |
| R | 0.0006** | 0.0004** | 0.0004** | 0.0047*** | 0.0013*** | 0.0014*** | 0.0014** | 0.0000ns | 0.0025*** | 0000ns | 0000ns | 0.0001. |
| B(R) | 0.0000ns | 0.0000ns | 0.0000ns | 0.0005ns | 0.0002ns | 0.0001ns | 0000ns | 0.0000ns | 0.0001ns | 0000ns | 0000ns | 0000ns |
| L | – | – | – | – | – | – | – | – | – | 0.1596*** | 0.1042*** | 0.1051*** |
| Y | 0.0153*** | 0.0125*** | 0.0065*** | 0.0789*** | 0.0391*** | 0.0370*** | 0000ns | 0.0001ns | 0.0001*** | 0.0165*** | 0.0094*** | 0.0069*** |
| G:L | – | – | – | – | – | – | – | – | – | 0.0206*** | 0.0126*** | 0.0302*** |
| G:Y | 0.0496*** | 0.0382*** | 0.0370*** | 0.1074*** | 0.0417*** | 0.0944*** | 0.0706*** | 0.0312*** | 0.0813*** | 0.0268*** | 0.0165*** | 0.0251*** |
| G:L:Y | – | – | – | – | – | – | – | – | – | 0.0628*** | 0.0281*** | 0.0529*** |
| ERROR | 0.0278 | 0.0202 | 0.0188 | 0.0475 | 0.0394 | 0.0252 | 0.0738 | 0.0461 | 0.0732 | 0.0520 | 0.0358 | 0.0404 |

<sup>a</sup> Variance represents proportion of the phenotypic values for Severity, Field Response and Coefficient of infection explained by the respective sources of variation
SEV, disease severity; RES, Field Response; CI, Coefficient of Infection (i. e. CI = SEV x RES).
G, genotype variance estimate; R, replication variance estimate; B(R), Block nested with in replication variance estimate; L, location variance estimate; Y, year variance estimate; G:L, genotype x location variance estimate; G:Y, genotype x year variance estimate; G:L:Y, genotype x Location x year variance estimate.
ns, not significant. $P < 0.1$ ; $*P < 0.05$ ; $**P < 0.01$ ; $***P < 0.001$

Next, we calculated the correlation coefficients to see the relationships between the three phenotypic variables within and across environments (Table 2). Correlation coefficient (*r*) for SEV of 2015 & 2016 data was 0.47 at Chefe-Donsa, 0.24 at Kulumsa and 0.70 at Meraro locations. Likewise, for RES, it was 0.43 at Chefe-Donsa, 0.33 at Kulumsa and 0.72 at Meraro. Almost a similar pattern of correlations for CI (0.51, 0.26 & 0.68) were observed at Chefe-Donsa, Kuluma and Meraro respectively (Table 2). This obviously shows that the level of the disease prevalence between the two years is different within the three location with a little bit of similarity for Merao test location. This again confirms the difference in reaction distribution of the genotypes as presented in Fig 1. Correlations among the three reaction data types was also assessed at different combinations (SEV vs RES, SEV vs CI & RES vs CI). High correlation was observed for SEV vs RES (*r* = 0.94 ± 0.01), SEV vs CI (*r* = 0.97 ± 0.01) and RES vs CI (*r* = 0.92 ± 0.01) within same environment while very low and nearly similar for within same location-different year and among locations (Table 2). This is expected because the level of disease pressure that result in a high severity is highly likely to result in higher reaction of the genotype exposed and hence positively impact the coefficient of correlation. Besides, using the T-distribution test for most of the comparisons shows, statistically significant correlations at *P < 5%* except very few marked as “ns” (Table 2). This can be partly explained by the higher degrees of freedom in the analysis (i.e. DF= 293 −1=291) due to the large number of samples.

**Table 2.** Ccorrelation coefficients of SEV, RES and CI values within and among six environments.

| SEV vs. SEV | CHD_15 | CHD_16 | KUL_15 | KUL_16 | MER_15 | MER_16 |
| --- | --- | --- | --- | --- | --- | --- |
| CHD_15 | 1 |  |  |  |  |  |
| CHD_16 | 0.47 | 1 |  |  |  |  |
| KUL_15 | 0.05<sup>ns</sup> | 0.30 | 1 |  |  |  |
| KUL_16 | 0.39 | 0.79 | 0.24 | 1 |  |  |
| MER_15 | 0.28 | 0.53 | 0.17 | 0.66 | 1 |  |
| MER_16 | 0.27 | 0.59 | 0.19 | 0.78 | 0.70 | 1 |

| RES vs. RES | CHD_15 | CHD_16 | KUL_15 | KUL_16 | MER_15 | MER_16 |
| --- | --- | --- | --- | --- | --- | --- |
| CHD_15 | 1 |  |  |  |  |  |
| CHD_16 | **0.43** | 1 |  |  |  |  |
| KUL_15 | 0.12 | 0.30 | 1 |  |  |  |
| KUL_16 | 0.38 | 0.76 | **0.33** | 1 |  |  |
| MER_15 | 0.30 | 0.54 | 0.23 | 0.66 | 1 |  |
| MER_16 | 0.30 | 0.56 | 0.22 | **0.77** | **0.72** | 1 |
| CI vs. CI | CHD_15 | CHD_16 | KUL_15 | KUL_16 | MER_15 | MER_16 |
| CHD_15 | 1 |  |  |  |  |  |
| CHD_16 | **0.51** | 1 |  |  |  |  |
| KUL_15 | 0.03<sup>ns</sup> | 0.31 | 1 |  |  |  |
| KUL_16 | 0.46 | **0.80** | **0.26** | 1 |  |  |
| MER_15 | 0.32 | 0.52 | 0.18 | 0.67 | 1 |  |
| MER_16 | 0.31 | 0.57 | 0.18 | 0.78 | **0.68** | 1 |
| SEV vs. RES | CHD_15 | CHD_16 | KUL_15 | KUL_16 | MER_15 | MER_16 |
| CHD_15 | **0.93** | **0.41** | 0.07 | 0.40 | 0.31 | 0.30 |
| CHD_16 | **0.49** | **0.96** | 0.33 | 0.78 | 0.54 | 0.57 |
| KUL_15 | 0.10<sup>ns</sup> | 0.28 | **0.91** | **0.28** | 0.19 | 0.19 |
| KUL_16 | 0.39 | 0.76 | **0.30** | **0.96** | 0.67 | 0.74 |
| MER_15 | 0.28 | 0.53 | 0.20 | 0.64 | **0.91** | **0.68** |
| MER_16 | 0.28 | 0.58 | 0.21 | 0.78 | **0.73** | **0.95** |
| SEV vs. CI | CHD_15 | CHD_16 | KUL_15 | KUL_16 | MER_15 | MER_16 |
| CHD_15 | **0.99** | **0.51** | 0.03 | 0.45 | 0.32 | 0.30 |
| CHD_16 | **0.48** | **0.96** | 0.28 | 0.84 | 0.58 | 0.62 |
| KUL_15 | 0.06<sup>ns</sup> | 0.32 | **0.95** | **0.26** | 0.18 | 0.20 |
| KUL_16 | 0.40 | 0.72 | **0.23** | **0.96** | 0.71 | 0.80 |
| MER_15 | 0.28 | 0.47 | 0.17 | 0.61 | **0.98** | **0.66** |
| MER_16 | 0.28 | 0.53 | 0.17 | 0.73 | **0.71** | **0.97** |
| RES vs. CI | CHD_15 | CHD_16 | KUL_15 | KUL_16 | MER_15 | MER_16 |
| CHD_15 | **0.92** | **0.53** | 0.07 | 0.44 | 0.32 | 0.32 |
| CHD_16 | **0.42** | **0.94** | 0.27 | 0.81 | 0.57 | 0.61 |
| KUL_15 | 0.07<sup>ns</sup> | 0.34 | **0.88** | **0.33** | 0.23 | 0.23 |
| KUL_16 | 0.41 | 0.73 | **0.27** | **0.94** | 0.68 | 0.81 |
| MER_15 | 0.31 | 0.49 | 0.19 | 0.62 | **0.91** | **0.69** |
| MER_16 | 0.30 | 0.53 | 0.17 | 0.72 | **0.68** | **0.95** |

Values in grey highlight represent same location but different years comparisons while bold face fonts indicate comparison within same locations and environments. Correlation coefficient values in bold face are the lowest and highest values obtained for the respective comparisons.
<sup>ns</sup> all Correlation coefficient values having this superscript represents statistically non-significant correlations between the tested data based on T-distribution

**Table 3.** Polymorphism of SNPs in Ethiopian durum wheat germplasm obtained from genotyping with Breeders’ 35K Axiom Array.

| Chr. | No. of SNPs | No Call Rate (Mean) | MAF |  | Heterozygosity |  | PIC |  | No of SNPs with MAF > 5%<sup>c</sup> |
| --- | --- | --- | --- | --- | --- | --- | --- | --- | --- |
|  |  |  | Mean | Range | Mean | Range | Mean | Range |  |
| 1A | 548 | 0.0034 | 0.2000 | 0.0034 - 0.5000 | 0.2766 | 0.0068 - 0.5000 | 0.2265 | 0.0068 - 0.3750 | 462 |
| 1B | 852 | 0.0038 | 0.2171 | 0.0017 - 0.5000 | 0.2879 | 0.0034 - 0.5000 | 0.2326 | 0.0034 - 0.3750 | 700 |
| 2A | 615 | 0.0037 | 0.1631 | 0.0017 - 0.5000 | 0.2399 | 0.0034 - 0.5000 | 0.2003 | 0.0034 - 0.3750 | 491 |
| 2B | 882 | 0.0037 | 0.1654 | 0.0017 - 0.5000 | 0.2380 | 0.0034 - 0.5000 | 0.1978 | 0.0034 - 0.3750 | 682 |
| 3A | 481 | 0.0039 | 0.2015 | 0.0034 - 0.5000 | 0.2768 | 0.0068 - 0.5000 | 0.2257 | 0.0068 - 0.3750 | 396 |
| 3B | 744 | 0.0038 | 0.1494 | 0.0034 - 0.5000 | 0.2214 | 0.0068 - 0.5000 | 0.1852 | 0.0068 - 0.3750 | 545 |
| 4A | 387 | 0.0040 | 0.1902 | 0.0034 - 0.4914 | 0.2708 | 0.0068 - 0.4999 | 0.2222 | 0.0068 - 0.3749 | 317 |
| 4B | 403 | 0.0037 | 0.1924 | 0.0034 - 0.4931 | 0.2737 | 0.0068 - 0.4999 | 0.2234 | 0.0068 - 0.3750 | 317 |
| 5A | 615 | 0.0038 | 0.1869 | 0.0034 - 0.4966 | 0.2663 | 0.0068 - 0.5000 | 0.2185 | 0.0068 - 0.3750 | 494 |
| 5B | 766 | 0.0037 | 0.2096 | 0.0034 - 0.5000 | 0.2930 | 0.0068 - 0.5000 | 0.2389 | 0.0068 - 0.3750 | 673 |
| 6A | 515 | 0.0036 | 0.2032 | 0.0034 - 0.4983 | 0.2782 | 0.0068 - 0.5000 | 0.2269 | 0.0068 - 0.3750 | 412 |
| 6B | 740 | 0.0040 | 0.1860 | 0.0034 - 0.4966 | 0.2664 | 0.0068 - 0.5000 | 0.2191 | 0.0068 - 0.3750 | 617 |
| 7A | 692 | 0.0043 | 0.1670 | 0.0034 - 0.4983 | 0.2410 | 0.0068 - 0.5000 | 0.2001 | 0.0068 - 0.3750 | 523 |
| 7B | 557 | 0.0040 | 0.1925 | 0.0034 - 0.4983 | 0.2701 | 0.0068 - 0.5000 | 0.2216 | 0.0068 - 0.3750 | 464 |
| Unknown<sup>a</sup> | 521 | 0.0059 | 0.1736 | 0.0017 - 0.4983 | 0.2486 | 0.0034 - 0.5000 | 0.2057 | 0.0034 - 0.3750 | - |
| NS match<sup>b</sup> | 823 | 0.0052 | 0.1815 | 0.0034 - 0.4983 | 0.2578 | 0.0068 - 0.5000 | 0.2125 | 0.0068 - 0.3750 | - |
| A genome | 3853 | 0.0038 | 0.1857 | 0.0017 - 0.5000 | 0.2623 | 0.0034 - 0.5000 | 0.2158 | 0.0034 - 0.3750 | 3095 |
| B genome | 4944 | 0.0038 | 0.1871 | 0.0017 - 0.5000 | 0.2634 | 0.0034 - 0.5000 | 0.2162 | 0.0034 - 0.3750 | 3998 |
| AB whole genome | 8797 | 0.0038 | 0.1865 | 0.0017 - 0.5000 | 0.2629 | 0.0034 - 0.5000 | 0.2161 | 0.0034 - 0.3750 | 7093 |

<sup>a</sup>Unknown, are SNPs not assigned to any of the chromosomes and has no position values in base pair as well. <sup>b</sup>These are SNPs with no significant match in the genome <sup>c</sup>Are final set of SNPs defined based on MAF > 5% and used for association and all other downstream analysis

### Genotypes/SNPs Called and Chromosomal Distribution

Genotype data assessment on the 300 accessions resulted in 296 which passed Dish Quality Control (DQC) ≥ 0.8. Heterozygosity and SNP analysis defined a final panel of 293 accessions with 10622 (30.39% of the 35K SNP array) polymorphic SNP markers (Fig 3) after excluding Individuals with Heterozygosity values of ≥ 3%. We extracted the physical positions (from RefSeqv1.0) of the informative SNPs which led to 3853 (36.27%) markers being assigned to the A-genome, 4944 (46.54%) to the B-genome, 481 (4.53%) to the D-genome and 521 (4.90%) to “Chromosome U” (i.e. unknown physical position); 823 (7.75%) markers had no significant match in the genome. Excluding the D-genome SNPs (as they are not expected in a tetraploid/durum wheat) the unknown set and those which had no significant match resulted in a total of 8797 physically positioned SNPs.

**Fig 3.**
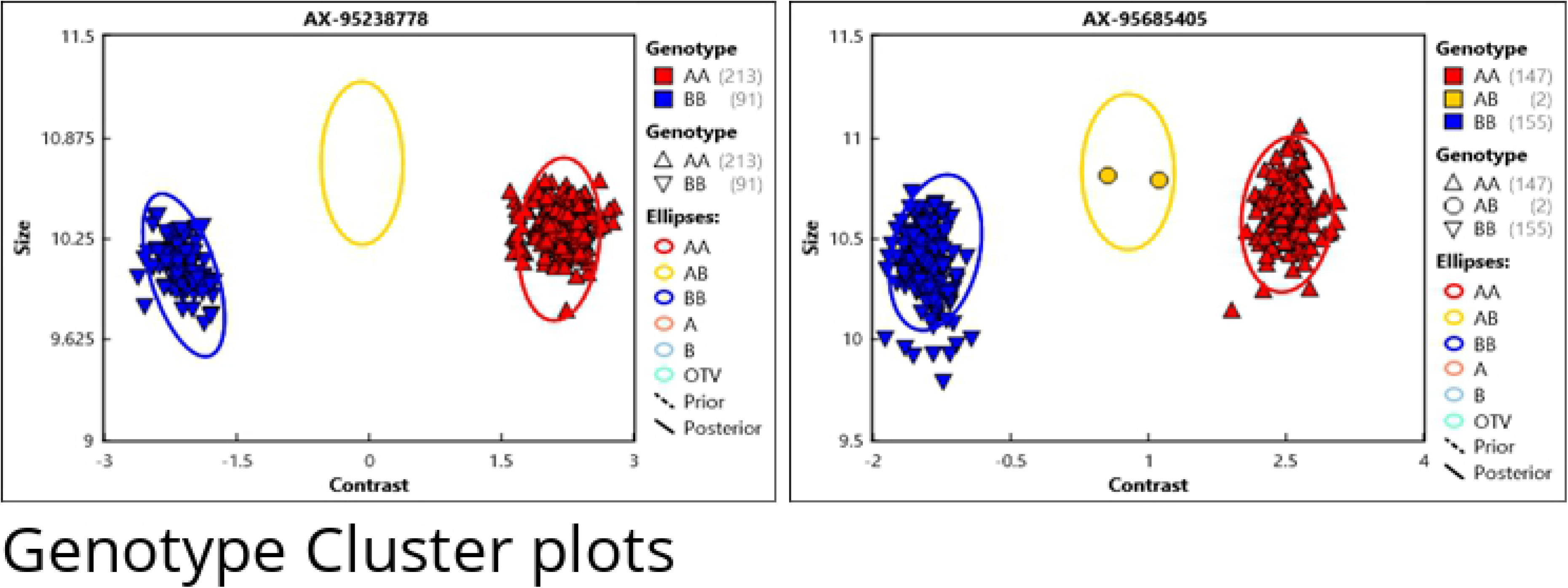
Examples of Cluster plots of durum association panel resulted from Axiom Analysis Suit V.2 with Best Practice Workflow. QC Threshold passed accessions are 296 and 8 samples are control; **A** & **B**are accessions genotyped with Axiom SNP probes AX-95238778 and AX-95685405 respectively

Varying numbers of SNPs, minor allele frequencies (MAF), Locus heterozygosity and polymorphic information content (PIC) value of the SNPs were obtained at chromosome, sub-genome, and whole genome levels (Table 4). The highest number of SNPs (852) were observed in chromosome 1B while the lowest (387) was found in chromosome 4A. Mean MAF was the highest (0.2171) for chromosome 1B while it was the lowest (0.1631) for chromosome 2A. Mean locus heterozygosity and PIC values were highest (0.2879 and 0.2326) for chromosome 1B SNPs and lowest (0.2214 and 0.1852) for chromosome 3B. In summary, we identified on average 628 ± 42 SNPs per chromosome.

**Table 4.** Linkage Disequilibrium estimate among Ethiopian durum wheat panel.

| Classes (Mbp) | Number of pairs | No of Significant Paris | % of Significant Paris | Mean $r^2$ | Mean of $r^2 > 0.2$ |
| --- | --- | --- | --- | --- | --- |
| 0 -10 | 9996 | 1367 | 13.68% | 0.28 | 0.36 |
| 10 -20 | 9979 | 795 | 7.97% | 0.16 | 0.29 |
| 20 - 30 | 9968 | 930 | 9.33% | 0.14 | 0.29 |
| 30 - 40 | 9956 | 996 | 10.00% | 0.13 | 0.29 |
| 40 - 50 | 9947 | 1146 | 11.52% | 0.13 | 0.29 |
| 50 - 60 | 9896 | 1317 | 13.31% | 0.13 | 0.29 |
| 60 - 70 | 9880 | 1355 | 13.71% | 0.13 | 0.30 |
| >70 | 558264 | 168665 | 30.21% | 0.11 | 0.34 |
| Total | 627886 | 176571 | 28.12% | 0.11 | 0.34 |

\*significance in LD was declared at $P \leq 0.001$ where $P$ represents probability of Disequilibrium (pDiseq)

We further explored the SNPS by looking at the sub-genome distribution. We found lower number of A genome (43.8%) than B genome (56.2%) SNPs (Fig 4). SNPs from both genomes had similar MAF (0.1857, 0.1871), locus heterozygosity (0.2623,0.2634) and PIC (0.2158, 0. 2162). Group 1 chromosomes had the highest SNP density/coverage and the most diverse and informative SNPs. For more reliable result from downstream analyses, these SNPs were further refined to maintain only those with a > 5% MAF cut-off resulting in a final set of 7093 SNPs (Table 3). The distribution of this final SNP set was on average 442.1 ± 27.0 for the A-genome and 571.1 ± 53.0 for the B-genome.

**Fig 4.**
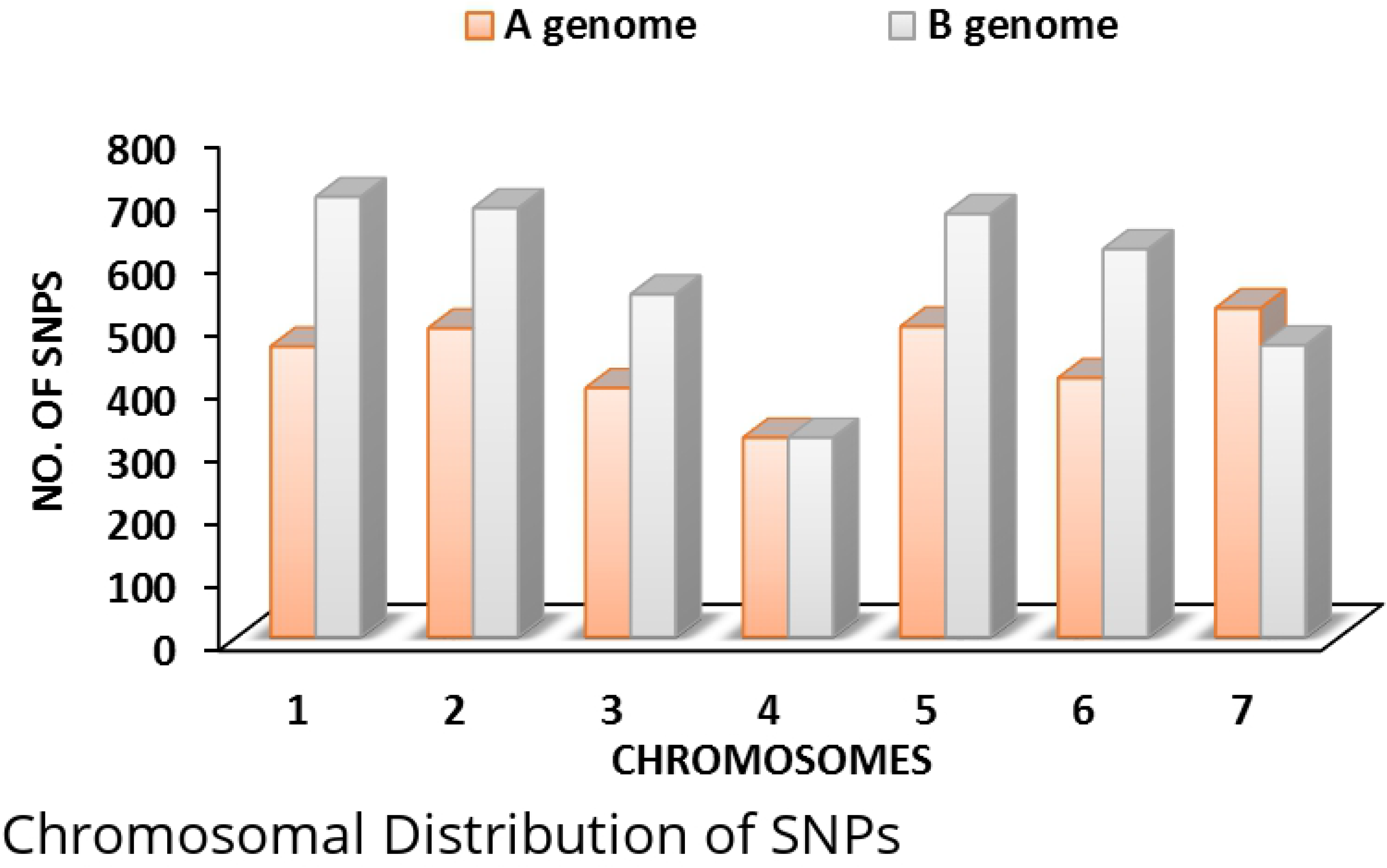
Chromosomal Distribution of 7093 SNPs across genome A and B. They are a subset of the 8797 SNPs (Table 4) and are selected for association analysis based on their MAF values of > 5%.

### Population Groups and Relatedness

We performed a population structure analysis which grouped the panel into two clusters. Landraces clustered together into one group while all the cultivars clustered in a second group (Fig 5A). Structure harvester analysis suggested that delta K attained its highest value at K=2 which is the most likely number of populations in the study panel (Fig 5B). In a global view, kinship analysis resolved the panel into two distinct groups as well, where the cultivars still stood out separately from the rest of the panel (Fig 5C). A closer look at the grouping pattern however revealed the presence of four groups where the cultivars (CL) clustered separately while the landraces separated into three sub-groups (LR-I, LR-II & LR-III). Principal Component Analysis (PCA) also resolved the panel into four clusters with the cultivars noticeably still isolated from the landraces (Fig 5D). Despite PCA revealed four clusters, only the first two components explained 66.67% (Fig 5E) of the variation which is in agreement with presence of two main groups as depicted by structure and kinship analyses.

**Fig 5.**
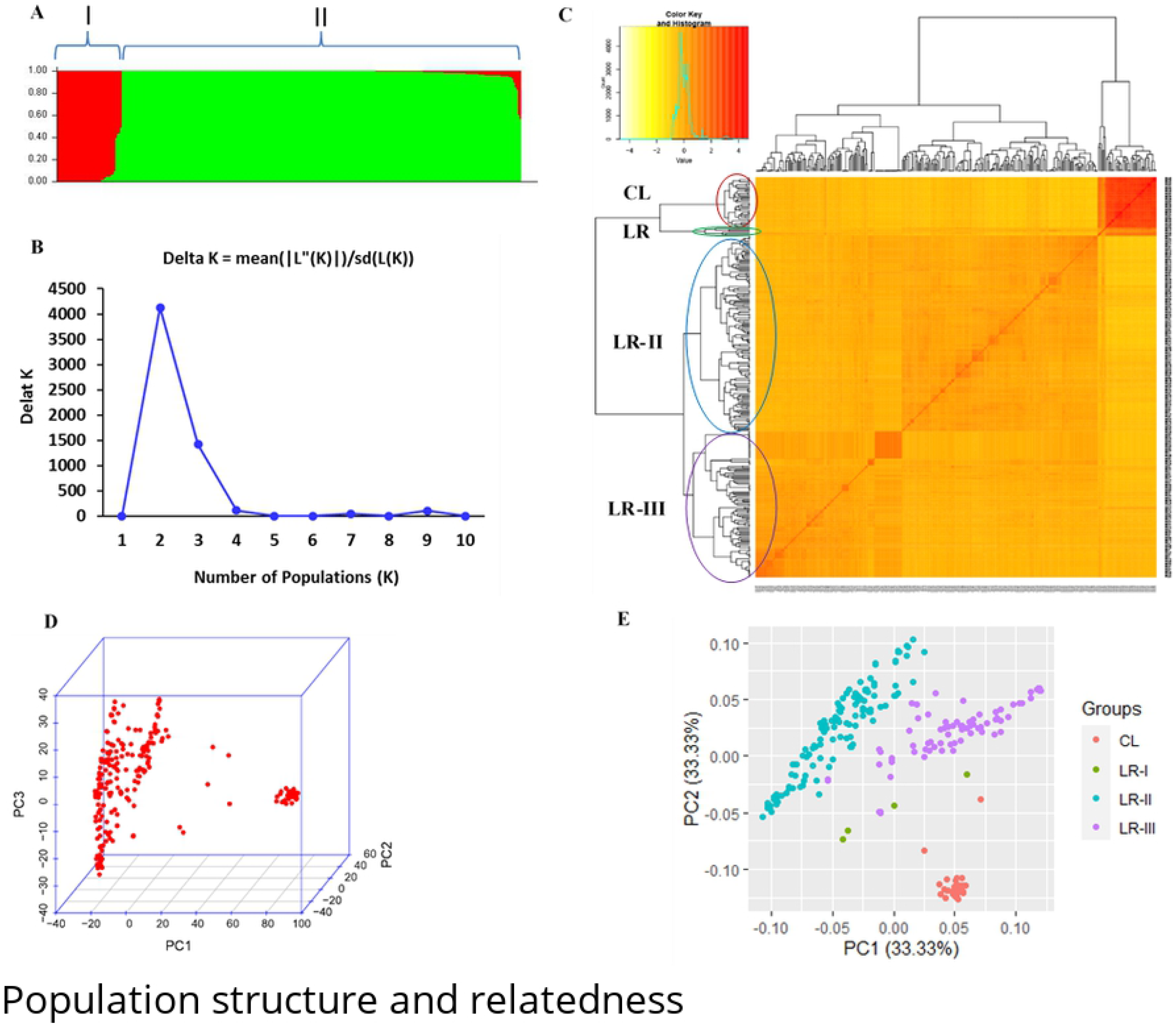
Population Structure and relatedness among the 293 accessions used for the GWAS. **A**) population structure plot as reveled by STRUCTURE analysis. **B**) Delta K plot from STRUCTRUE HARVESTER analysis. **C)** 293×293 Kinship matrix plot using genetic similarity matching where outside the matrix is a clustering tree of the panel in to 2 main groups (in global view) while a bit of detailed view gave four clusters: CL= Cultivars and three Land Races sub-groups (LR-I, LR-II and LR-III). At the top left corner of the plot is the distribution of estimated kinship values. **D)** PCA plot of the first three principal components where improved varieties still stood out in a distinct group at the very right end in the plot while the rest grouped in a similar way to the Kinship plot. **E)** 2D plot elaborating sub-groups as identified by the PCA analysis.

### Linkage Disequilibrium

We computed the squared allele frequency correlation (r^2^) for all pairwise comparisons of distances between SNPs using TASSEL. At a genome-wide level, 627,886 inter-SNP distances were found through binning LD analysis output at a bin of 1000 bp. Of this, 176571 (28.12%) were in significant LD with an average LD estimate of r^2 = 0.11 where the highest value (0.28) was achieved within the first 10 Mbp of physical distance (Table 4). The mean LD above the critical value (r^2 = 0.2), was also the highest (0.36) within the same 10 Mbp of physical distance as reported for the total (Table 4).

The fitted LOESS curve intersected with the critical LD value at physical distance of 69.1 Mbp where all the values of LD bellow this point were considered to be due to physical linkage among the intra-chromosomal loci/SNP pairs (Fig 6). The LD started to decay below this critical value to an average r2 of 0.16 for an increase of 10 Mbp (in the interval 10-20) suggesting that the overall LD accounted for the association is exhibited by relatively a shorter genomic distance.

**Fig 6.**
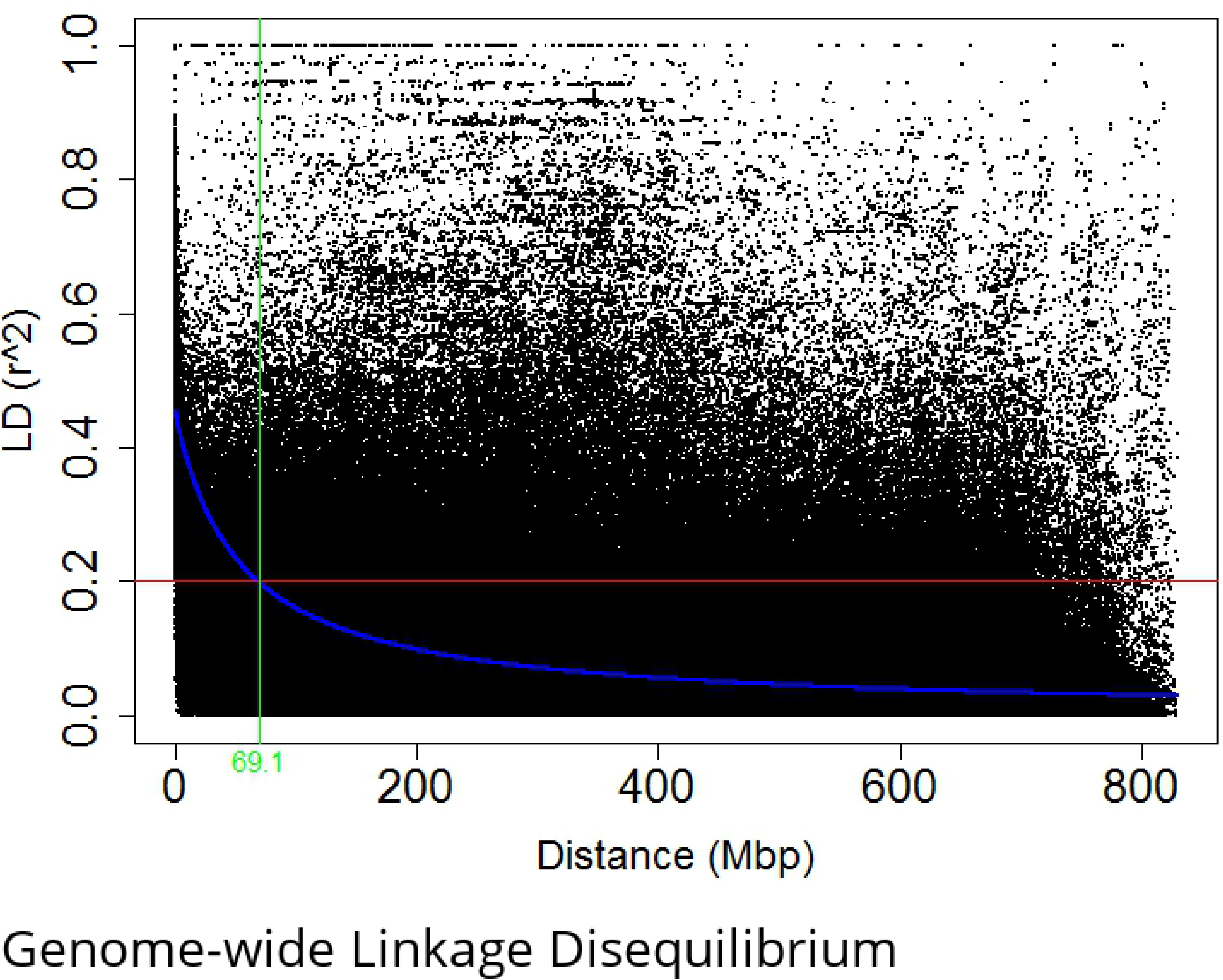
Genome-wide Linkage Disequilibrium as dictated by physical distance. Average pair-wise inter-SNP LD (*r^2^*) values plotted against physical distance in base pairs based on the wheat reference genome RefSeq v.1.0. The red line indicates the threshold LD.

### Marker-Trait Associations for Resistance to Yellow Rust

The association analysis of CI resulted in a total of 12 SNPs, across four chromosomes (1A, 1B, 2B, and 7B), significantly associated with yellow rust resistance at FDR-adjusted *P≤0.05* (Table 5). Five SNPs (AX-94534607, AX-95115308, AX-94482796, AX-94856684, AX-94827580) at CHD and one SNP (AX-94648330) at KUL were identified as location specific resistance associated SNPs. Six of the SNPs (AX-95171339, AX-94436448, AX-95238778, AX-95096041, AX-94730403 & AX-94427201) however, were consistently identified (Table 5 & Fig 7) at each location and combined analysis across all six environments (BLUE-all). No location specific SNP was identified at MER. Almost similar sets of SNPs were identified from GWAS analysis for SEV and RES as well. As both SEV and RES are highly correlated to each other and to CI (Table 2), we here focused reporting the GWAS results in relation to CI. A comprehensive result of the GWAS analysis for SEV, RES and CI of each environment data is presented in Table S2. The phenotypic variation R^2^ explained by all six SNPs on chromosome 1B ranged from 51.8 – 54.1% for CHD, 47.8 – 54.1% for KUL and 59.6 – 61.0% for MER (Table 5). MAF ranged from 14.2% to 31.7% with the highest for AX-94534607, AX-94856684 and the lowest for AX-95171339. The lowest (5.70E-09) significant FDR_Adjusted_P-value was exhibited by AX-94730403 on chromosome 1B. The resistance allele frequency (RAF) ranged from 0.20 to 0.83 among the panel with marker AX-94730403 (on chromosome 1B) attaining the lowest value while AX-95115308 (on chromosome 1A) having the highest value.

**Table 5.**
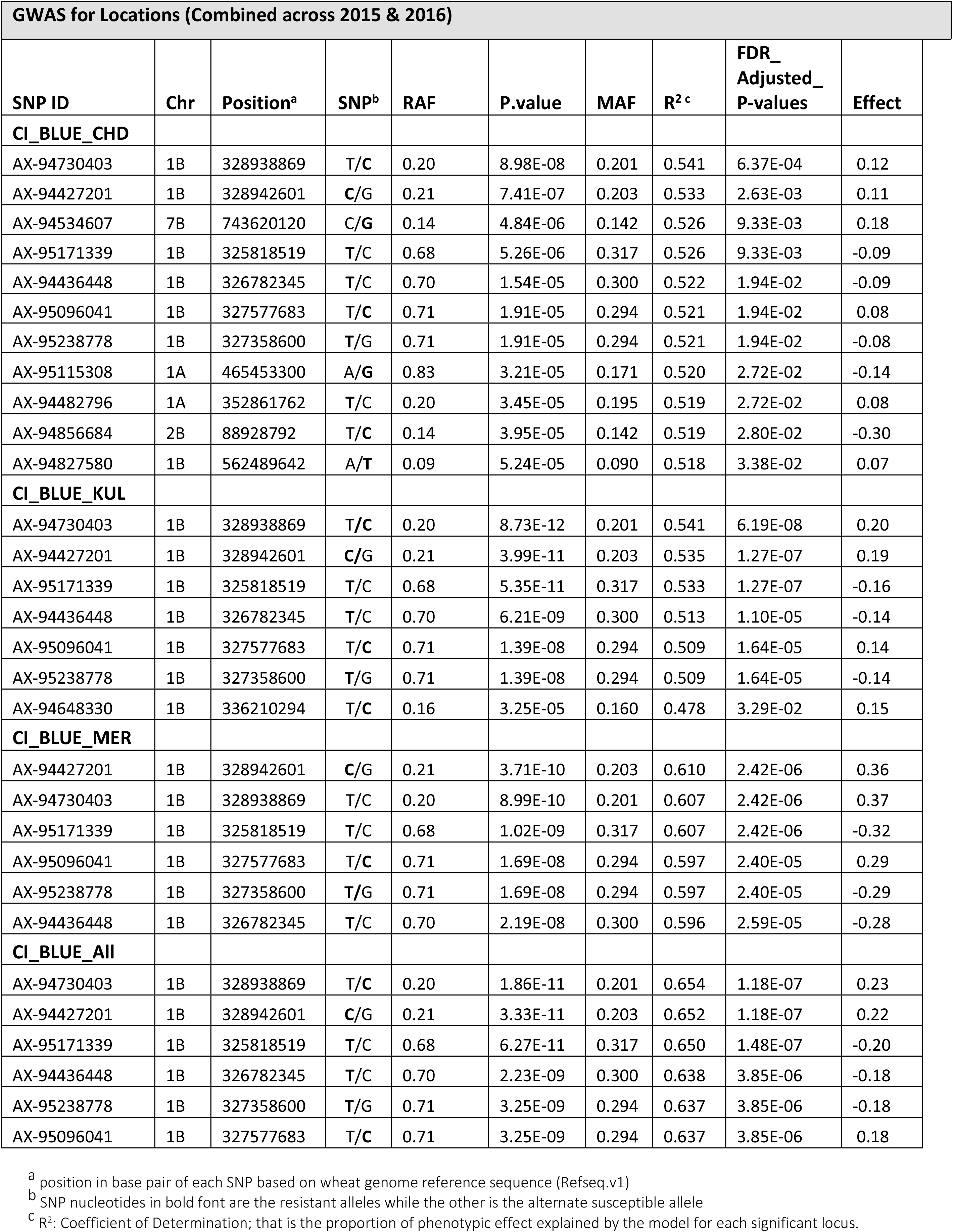
Markers significantly associated with Yellow rust resistance identified by GWAS analysis.

**Table 6.**
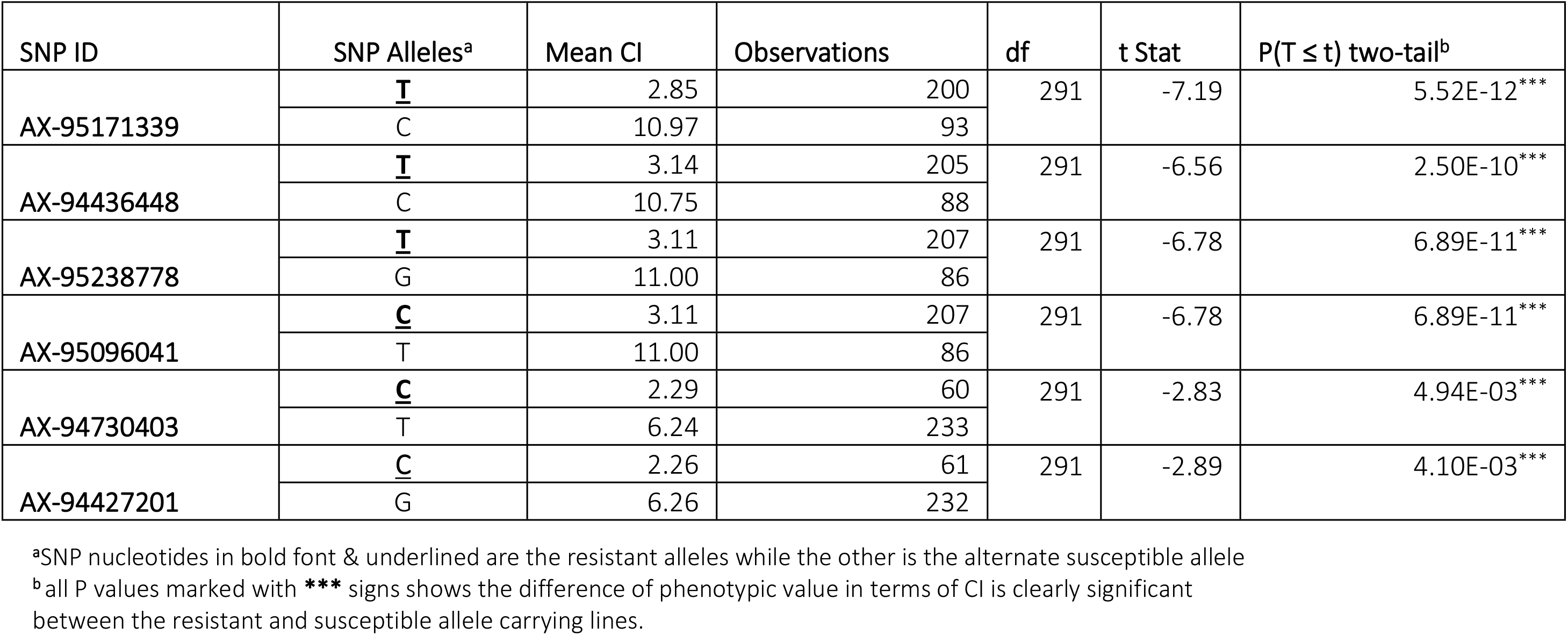
Single locus-based variation of phenotypic values of resistant and susceptible alleles as resulted from Two-Sample T-test Assuming Equal Variances.

**Fig 7.**
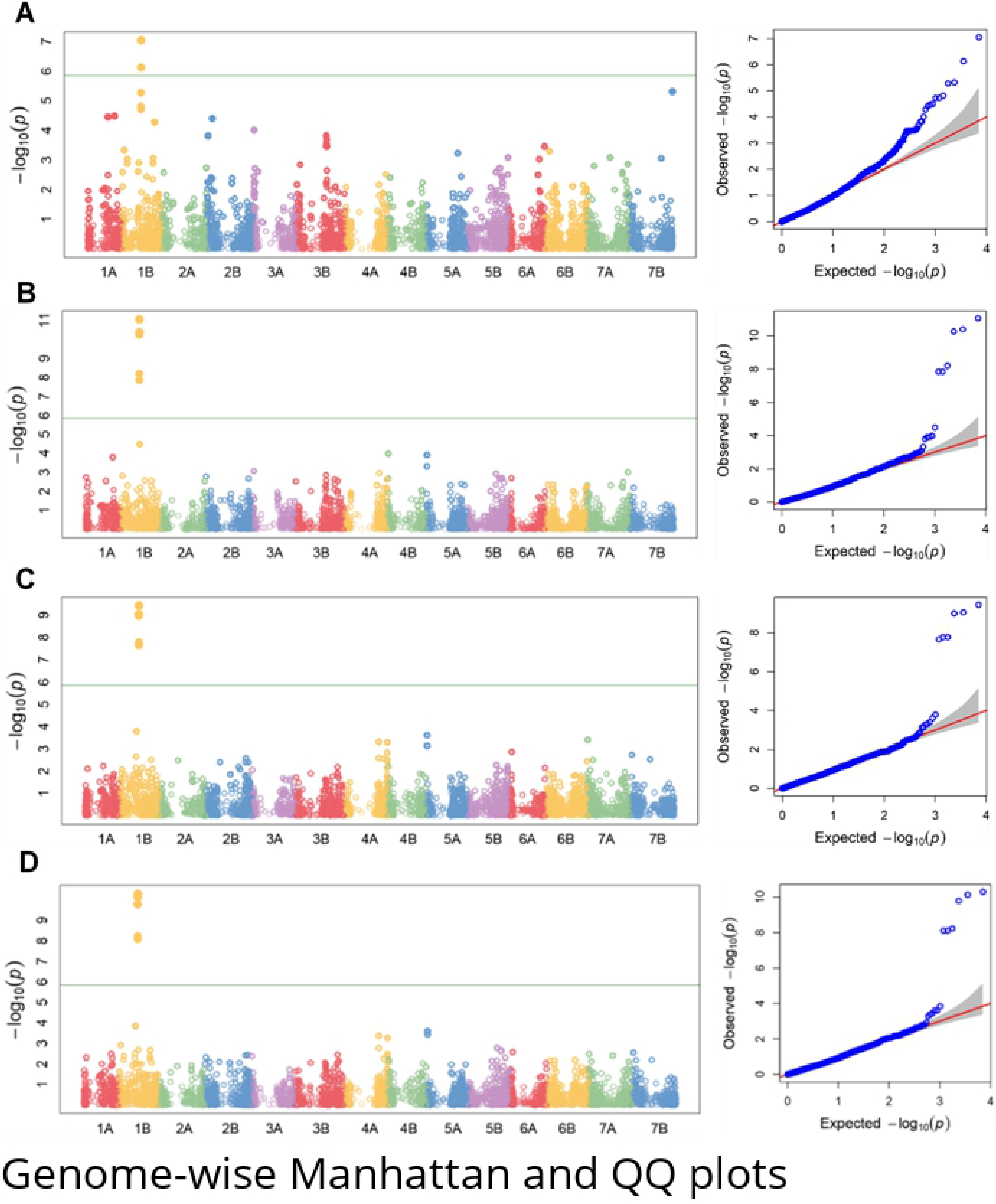
Genome-wise Manhattan and QQ plots of GWAS of yellow rust resistance in Ethiopian durum wheat. Results represent association analysis of CI data from 293 durum wheat accessions at Chefe-Donsa (A), Kulumsa (B) Meraro (C) and combined across the six (3 locations by 2 years) environments (D). Accessions were genotyped with Breeders' 35K Axiom Array for wheat.

### Other Genes in The Vicinity Of SNPs Associated With The Pst Resistance

Further investigation on approximately 4.15 Mbp (325818004-329960910) genomic region encompassing the identified resistance associated SNPs on chromosome 1B indicated the presence of 36 genes and protein families (Table 7). Four of these genes (MATH domain containing protein (TraesCS1B01G180400), Alpha-galactosidase (TraesCS1B01G181700), Chloroplast inner envelope protein putative, expressed (TraesCS1B01G182300) and Plant basic secretory family protein (TraesCS1B01G182700)) were redundantly found. SNP AX-95171339-1B, which is found close to the outer most of this region is very closely associated to Pentatricopeptide repeat-containing protein (TraesCS1B01G179700) while SNP AX-94436448 is flanked by DNA-directed RNA polymerase subunit (TraesCS1B01G179900) and Peptide chain release factor 2 (TraesCS1B01G180000). The ABC transporter gene family (TraesCS1B01G181200) which is known to confer durable resistance to multiple fungal pathogens in wheat [71] is also very closely associated to AX-95096041. Similarly, the disease resistance protein RPM1 (TraesCS1B01G182900) belonging to known disease resistance protein families (NBS-LRR class) was also found proximal to SNP AX-94427201-1B.

**Table 7.**
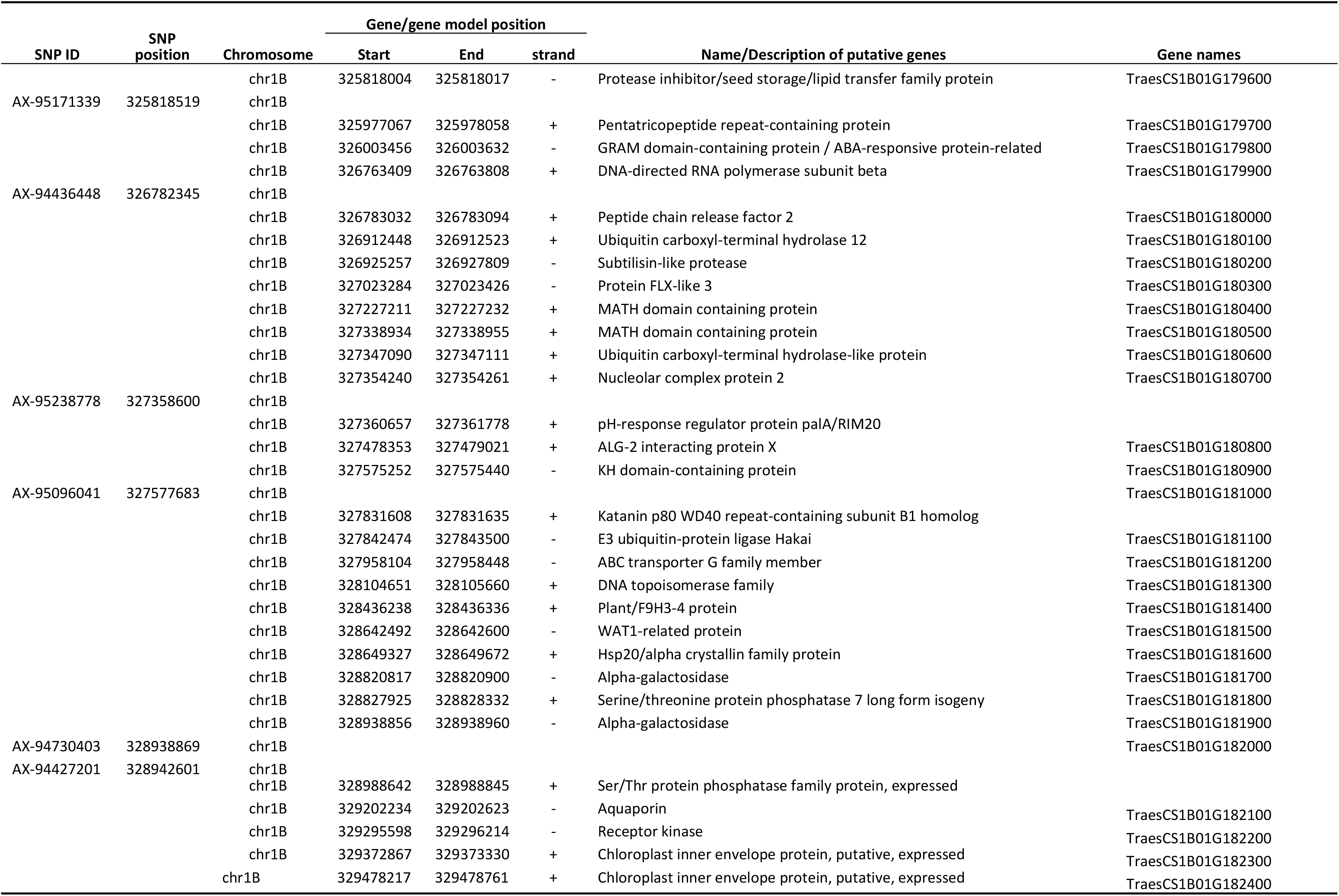
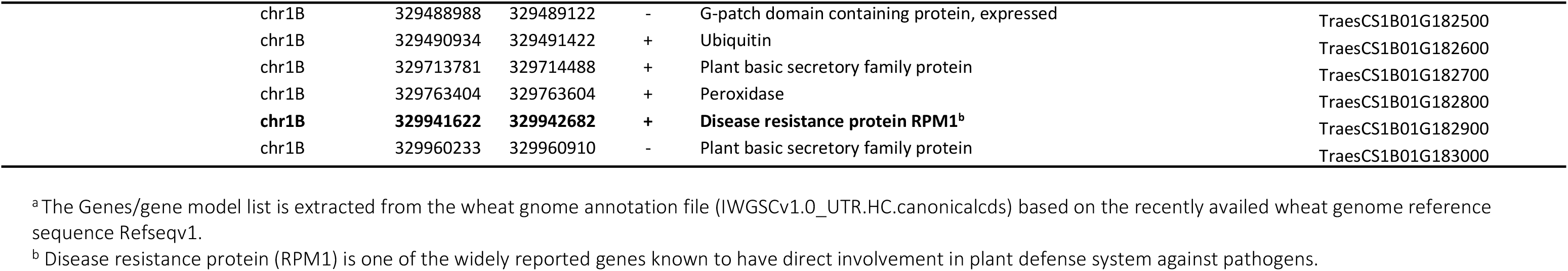
Genes and gene models^a^ reported with in the identified Yr resistance associated region and close to the nearest significant SNP.

## Discussion

### Phenotypic Variability in Resistance to Pst

The average field response to *Pst* of accessions was relatively low among the test sites. This is especially clear when examining the result of Chefe-Donsa and Kulumsa sites for which the mean SEV, RES and CI values are 1.4, 0.1, 1.1 and 3.6, 0.1, 2.5, respectively (Fig 1; Table 1). Notably, response at Meraro was higher because the location is known to be one of the hot spot sites for *Pst* infestation, hence why it is usually used as a stripe rust test site. On the other hand, the average SEV (5.33%), RES (0.24), CI (3.92) in 2015 is lower than it was in 2016 SEV (8.90%), RES (0.23) and CI(6.94) which might be attributed to a shortage of moisture in 2015. It is known that establishment of *Pst* infection in the field is highly dependent on available moisture and cool night temperatures which ultimately affects disease development. Very few accessions (18) consistently appeared as resistant across all the three locations which could be due to the broad-spectrum resistance of the accessions for the *Pst* race composition at the testing sites. Overall, the reaction data was unevenly distributed particularly for Kulumsa and Chefe-Donsa sites, although applying transformations to the SEV, CI and RES data improved the distribution (Fig 2D, E and F). Genotypic variances at CHD_2015 and KUL_2015 were non-significant which is somehow expected because of the low disease pressure as discussed above. Variance component due to blocking nested within replication was not significant which indicates that variation due to nesting was negligible. Particularly, genotype by environment interactions have significantly varied at both location and combined level which most likely is due to variation in disease pressure among the locations. Relatively varying levels of correlations for the disease reaction data (SEV, RES & CI) were observed within locations indicating level of disease pressure-based responses of the genotypes. On the other hand, correlations for Inter-disease reaction data combinations (Table 2) among environments were very high, ranging from 0.91-0.96 for SEV vs RES; 0.95-0.99 for SEV vs CI and 0.91-0.95 for RES vs CI. This provided the basis for performing the GWAS analysis on any of the three disease reaction data although CI is preferred as it combines representation of both SEV and RES.

### Population Structure, Relatedness and LD

The population structure analysis plot clustered the panel into two distinct groups where members of group I are mainly (36/39) improved durum cultivars while that of group II contains all the landraces besides three cultivars. This was in agreement with the structure harvester output that suggests K=2 is the most likely grouping value of the panel. The presence of the two groups in the population structure and further subgrouping in the Kinship has not shown any significant correlation with the pattern of the phenotypic values (resistance reaction) among the panel. Consequently, the data led to identification of a true and acceptable marker-trait association for the resistance as opposed to the discovery of false positive association.

### Analysis of the major resistance locus identified on chromosome 1B

As expected, the significant genotypic variance among the panel was reflected in the presence of the identification of significant marker-trait associations at various level. Despite the significant difference found among location variance components, similar sets of SNP association were identified both for location and the combined analysis. This probably indicates that variation in disease pressure may not as such affect sets of identified significant associations rather it affects their strength. On the other hand, the five significant associated SNPs identified only at Chefe-Donsa could mean presence of *Pst* race specificity at this testing site relative to the others. The consistence occurrence of the six significantly associated SNPs probably has to do with a wide spectrum effectiveness of the resistance gene/genes underlying the loci as well. Year based combined association analysis for 2015 resulted in the identification of the same six SNPs at FDR-adjusted *P≤0.05* while at a very high level of significance for 2016 (S2 Table). Such weaker association in 2015 may partly be explained by the less disease pressure occurred in the year 2015 as compared to the higher severity in 2016 (S2 Table). Likewise, R^2^ for year based combined data ranged from 51.4-52.0% and 63.3%-65.7% for 2015 and 2016 respectively (S2 Table). This situation indicates the existence of more severe disease pressure in 2016 than in 2015 which indirectly confirms why SNPs identified in 2016 data are more highly significant than those identified in 2015.

Our data suggests that chromosome 1B is an important contributor of loci significantly associated with the resistance as eight of the twelve identified SNPs are located on it. Particularly in the BLUE_all GWAS, all six associated SNPs were from this chromosome. Several genes associated with yellow rust resistance in wheat have been reported on chromosome 1B from multiple GWAS studies. This highlights its usefulness and why efforts to further define these loci are warranted. So far, 84 *Pst* resistance genes have been designated [72] and many QTLs identified in wheat have been reported on chromosome 1B [73]. *Yr10* [74], *Yr9* [75], *YrAlp* [72]*, Yr15* [76] *, YrH52* [77]*, Yr64, Yr65* and *Yr24/Yr26* [78], *YrExp1* [72]*, Yr29/Lr46* [79] are some of the known YR genes identified on chromosome 1B and derived from wild relatives and cultivars. Interestingly, the recently cloned Yr15 gene is in position 547Mb on CS and the top markers identified in the current study are in position 325-330 Mb suggesting that the identified loci is different to Yr15. Two additional SNPs were identified on chromosome 1A while the other two came from chromosome 2B and chromosome 7B. Several yellow rust QTLs and Yr genes are mapped on these chromosomes including Yr5 on chromosome 2B [80] which could be amongst the few that are effective against Ethiopian Pst races [45, 81].

To the best of our information, this is the first GWAS study in Ethiopia durum wheat across the three test locations for identification of marker trait association (MTAs) for *Pst* resistance. However, a similar study conducted on Ethiopian durum wheat in the USA led to the identification of 12 loci associated with resistance to *Pst* on seven chromosomes of which chromosome 1B is one of them besides chromosomes 1A, 2BS, 3BL, 4AL, 4B and 5AL [17]. On the other hand, [82] carried out a GWAS on synthetic hexaploid wheat at Meraro and Arsi Robe and reported a total of 38 SNPs on 18 genomic regions associated with adult plant resistance. Some of these reported genomic regions are also identified on chromosome 1B besides 1A, 2B and 7B which is in agreement with the current study. So, similarities of this genomic regions in response to Pst resistance across various similar studies signifies the potential usefulness of the genomic regions for wheat resistance improvement.

## Conclusion

We identified a major genomic region on chromosome 1B harbouring six SNPs associated with resistance to *Pst* at adult plant stage consistently at each location and combined data. This suggests the presence of a gene or genes conferring resistance to *Pst* within this genomic region. The other associated SNPs identified only in one of the sites may highlight the presence of resistance gene/genes effective to location specific *Pst* races. This on the other hand calls for a separate consideration in future breeding strategies for durable Pst resistance enhancement in wheat. The study also identified effective sources of resistance to Ethiopian *Pst* races in Ethiopian durum wheat landraces that can be used, alongside the markers identified here, to transfer this locus into adapted cultivars to provide resistance against *Pst.* However, the diagnostic value of the identified SNPs needs to be further investigated to perfectly define the region and validate them in an independent germplasm. In general, the identified SNPs/resistance locus, coupled with the identification of multi-environment stable genotypes for resistance, will enhances the fight towards mitigation of *Pst* as it presents a double layer challenge both to the wide spectrum and site specific virulent *Pst* races.

## Acknowledgement

The authors are grateful to Kulumsa (KARC) and DebreZeit Agricultural Research Centre (DZARC) for kindly provision of experimental plot & cultivars; Ethiopian Biodiversity Institute (EBI) and EOSA (Ethio-Organic Seed Action) for their kind provision of the landrace accessions. This research was financially supported by the UK Biotechnology and Biological Sciences Research Council (BBSRC) under the Sustainable Crop Production Research for International Development (SCPRID) programme, the Ethiopian Institute of Agricultural Research (EIAR) and Addis Ababa University (AAU).

## Supporting Information

**S1 Table. Ethiopian durum wheat Landraces and Cultivars used for the GWAS**

**S2 Table. Association analysis result for all single and combined environment data**

